# Uncovering bioactive metabolites from the *Taxus wallichiana* endophyte *Annulohypoxylon purpureonitens* using reverse metabolomics

**DOI:** 10.64898/2026.08.17.744784

**Authors:** Tara Shrestha, Maria Rosario Garicia Gill, Heriberto Velez, Anjela Dahal, Samriddhi Bhochhibhoya, Santosh Lamichhane, Dhruva Prasad Gauchan

## Abstract

Endophytic fungi associated with the Himalayan yew (Taxus wallichiana) are an underexplored source of bioactive secondary metabolites. Here, we investigated the extracellular metabolites produced by *Annulohypoxylon purpureonitens*, an endophytic fungus isolated from T. wallichiana collected in Nepal, using bioactivity-guided screening coupled with liquid chromatography–tandem mass spectrometry. The fungal extract exhibited broad-spectrum antibacterial activity, with the strongest inhibition observed against Staphylococcus aureus and Enterococcus faecalis (minimum inhibitory concentration, 500 μg ml ¹). The extract also showed antioxidant activity and cytotoxic effects against HeLa and MCF-7 human cancer cell lines. Molecular networking based on the Global Natural Products Social Molecular Networking platform, together with reverse metabolomics, enabled the putative annotation of diverse metabolites, including hydroquinidine, chlorogenic acid, muramic acid and cordycepin conjugates. Several of these metabolite features were also observed across publicly available microbial metabolomics datasets. These findings highlight *A. purpureonitens* as a promising source of chemically diverse metabolites with antibacterial, antioxidant and cytotoxic activities and provide a basis for subsequent isolation and functional characterization of its bioactive constituents.

## Introduction

Endophytes that reside asymptomatically within plant tissues are increasingly recognized as prolific producers of bioactive compounds, often closely resembling those of their hosts [1]. Endophytic fungi, in particular, are a rich source of structurally diverse secondary metabolites that can outcompete pathogens for nutrients and/or induce host defence responses [2]. As key components of the plant microbiome, they contribute to host health, growth promotion, and resilience to abiotic and biotic stresses [3]. Over recent decades, endophytic fungi have been reported to produce numerous classes of bioactive secondary metabolites, including terpenoids, alkaloids, steroids, quinones, phenolics, and polyketides [4]. Many of these compounds exhibit antioxidant, antimicrobial, anticancer, and anti-inflammatory activities, underscoring endophytes as valuable biotechnological resources for natural product discovery [5].

The Himalayan yew (*Taxus wallichiana Zucc.)* is a high-value medicinal plant renowned for producing the anticancer drug paclitaxel (Taxol) [6]. In recent years, the plant-associated microbiota, particularly endophytic fungi, has attracted substantial interest as a potential source of bioactive metabolites. Among these endophytes, *Annulohypoxylon spp.* has emerged as promising candidates because they produce a spectrum of biologically active secondary metabolites [7]. *Annulohypoxylon* (family Xylariaceae) comprises ascomycetous fungi commonly occurring as saprobes on decaying wood and as endophytes in plant tissues. The focal species was originally described as *Hypoxylon purpureoniten* and was reassigned to *Annulohypoxylon* in 2005[8]. Members of this genus are taxonomically and chemically related to *Hypoxylon spp*. and are known to produce diverse secondary metabolites, including benzenoids, azaphilones, cytochalasins, and terphenyl derivatives [9]. Many of these compounds display potent bioactivities, such as antimicrobial, cytotoxic, and antioxidant effects [10]. For example,[11] reported that hypoxylonol F secreted by *Annulohypoxylon annulatum* shows potential for the treatment of diabetes mellitus. Metabolomics studies have further identified compounds such as methyl xylariate C, piliformic acid, and tensyuic acids that contribute to moderate-to-strong antibacterial activity [12]. Despite the chemical richness of xylariaceous fungi, the metabolomic landscape and bioactivities of *Annulohypoxylon sp*. remain insufficiently characterized. Addressing this gap is important for discovering novel metabolites and evaluating their pharmacological relevance. Among available analytical approaches, liquid chromatography–mass spectrometry (LC–MS) is particularly well suited for metabolomics due to its high resolution, sensitivity, and broad metabolite coverage.

Therefore, this study aims to (i) evaluate the antimicrobial, antioxidant, and anticancer activities of *A. purpureonitens* and (ii) perform comprehensive metabolite profiling using high-resolution LC–MS/MS to identify secondary metabolites with potential biological and pharmaceutical relevance, thereby prioritizing candidates for downstream functional and translational studies.

## Materials and Methods

### Sample collection and Isolation

The isolation of fungus was done from leaves of *Taxus wallichiana* collected from Taplejung (27°24.600N, 087°45.415E:2907m altitude), Nepal. The collected sample was sterilized following Gauchan et.al.,2021.The leaves segments were then inoculated onto Modified Melin-Norkrans medium (MMN) supplemented with chloramphenicol 0.05g/l to prevent contamination by endophytic bacteria[13]. Each plate was inoculated with four cut pieces of leaves. The plates inoculated were incubated at 27 ± 3°C for approximately 10 days. Sub-culturing was performed to acquire pure cultures. Morphological identification of the fungi was conducted using the original culture plates and microscopy was performed using Lacto Phenol cotton blue (LPCB)dye at 40x and 100x magnification[14].The resulting isolate was designated with the laboratory code T2B.

### Molecular Identification (DNA extraction and PCR)

Genomic DNA (gDNA) was isolated using Fungal/ Bacterial Mini-Prep Kit Quick DNA™ obtained from Zymo Research Company, with minor modifications[7]. The extracted gDNA was then amplified using PCR. The details of internal transcribed spacer (ITS) primers used and PCR conditions are given in Table 1. The obtained product was quantified using Qubit fluorometer in Dhulikhel Hospital, Nepal and sequenced using Sanger sequencing based on capillary electrophoresis (CES) method by Macrogen, Korea. The obtained sequence was trimmed and made consensus sequence was generated using CodonCode Aligner(version 12.0.1). The sequence was compared against the GenBank database using nucleotide BLAST. The phylogeny tree is constructed comparing our sequences with similar sequence from NCBI Genebank.

**Table 1.**
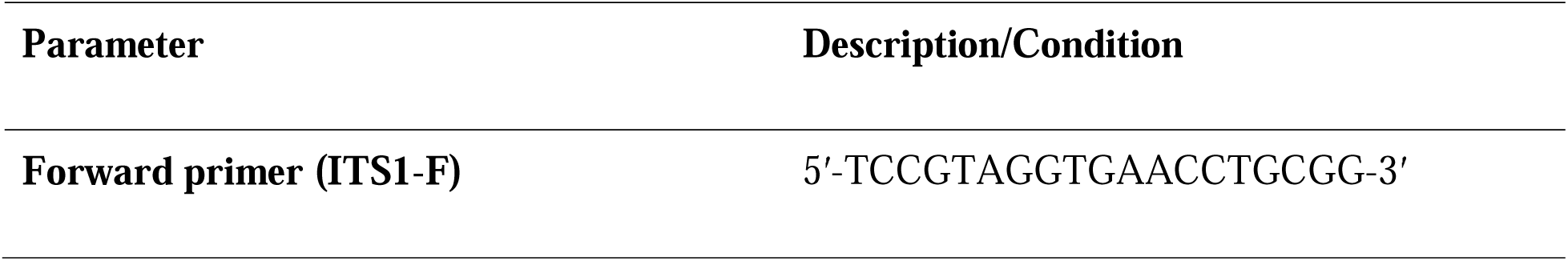
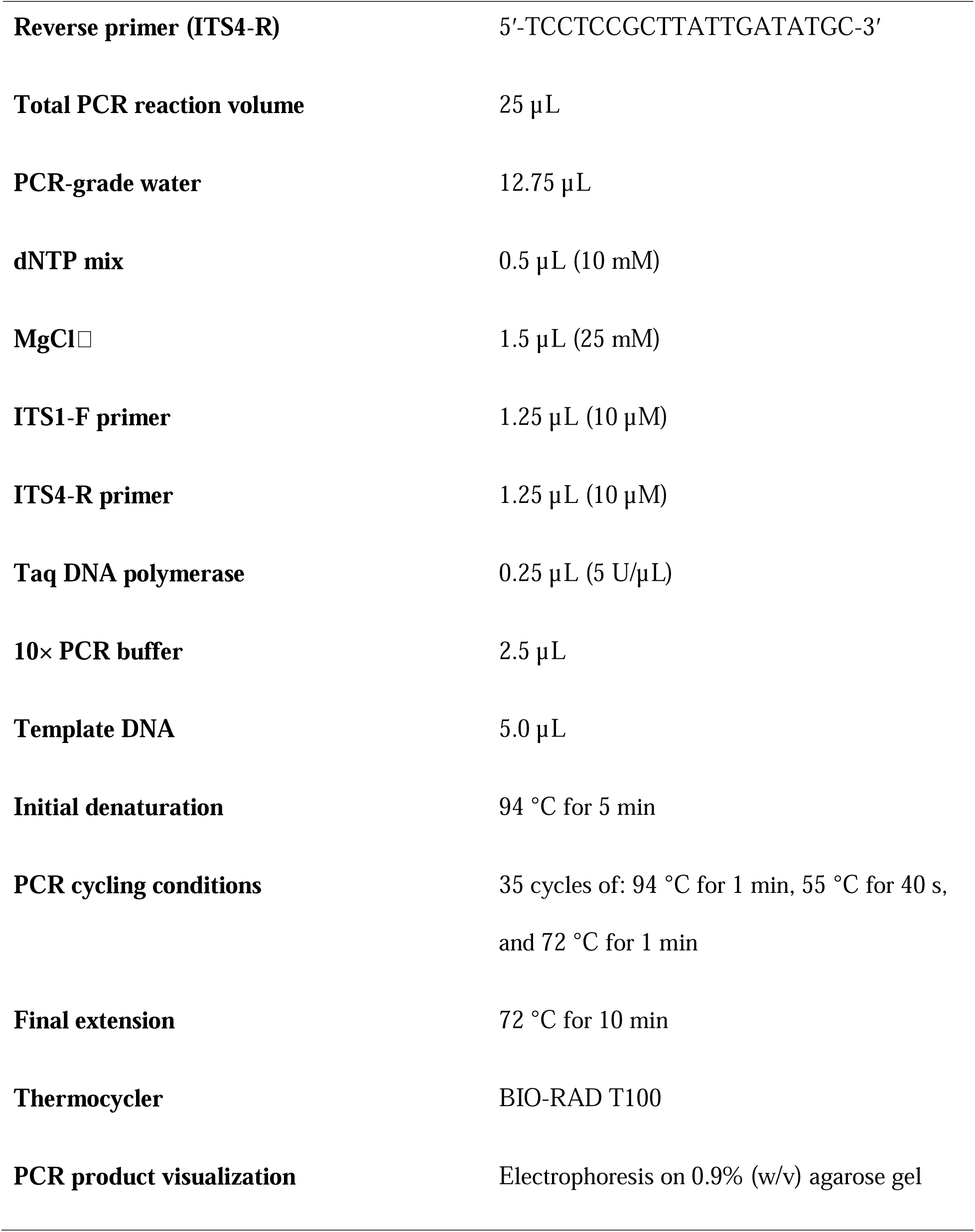
Primers used and PCR conditions employed for the amplification and sequencing of fungal DNA regions.

### Extraction of Fungal extract

The extraction was carried out for extracellular extract of *A. purpureonitens*. The mycelial plugs of 7mm each were plugged out from the pure culture and inoculated in a Erlenmeyer flask(250ml) with 100 mL of Potato Dextrose Broth (PDB) and kept at 150 rpm in a shaking incubator for about 2 weeks. After 2 weeks the mycelia and liquid filtrate were separated using a cheese cotton cloth. In liquid broth equal amount of ehtyl acetate was added, shaken well and separated using separating funnel. The filtrate was filtered using Whatman’s filter no., then solvent was evaporated using rotary evaporator. The obtained extract is store in 4^oC^ until further use [15, 16].

### Antimicrobial activity of Fungal Crude Extract

Antimicrobial activty of fungal extract was performed using agar well diffusion method against five ATCC(American Type Culture Collection) bactrial strains: two Gram-negative bacteria – *Klebsiella pneumoniae* (ATCC 13883) and *Escherichia coli* (ATCC 25922), two Gram-positive bacteria – *Staphylococcus aureus* (ATCC 12600) and *Enterococcus faecalis* (ATCC 19433), with one fungus *Candida albicans* (ATCC 10231)(Gauchan et al., 2021).The bacterial strains were initially cultured and grown overnight 37°C in Mueller-Hinton broth(MHB) whereas, *C. albicans* was grown in PDB at 27°C for 48 hours. All test strains were maintained to a McFarland turbidity standard of 0.5 McFarland standard (1× 10^8^ CFU/ml). Pathogens were spread on Mueller-Hinton (MH) agar allowed to dry for a few minutes. Once the plates dried, wells were created using a sterile borer with an inner diameter of 6mm. In one well, 20 µL of DMSO was added as a negative control. In the next well, 20 µL of fungal extract prepared in DMSO (25 mg/mL) was added. 10 µg Gentamicin (Ezy MIC strip discs, HIMEDIA) was used as a positive control for the bacterial plates, while for *C. albicans*, 24 µg of Fluconazole (Ezy MIC strip discs, HIMEDIA) was used. The plates were allowed for proper diffusion of extract and then incubated at 37°C for 24 hours, whereas plates with *C. albicans* were incubated for 48 hours at 27°C. The assay for each test pathogen was carried out in triplicate.The zones of inhibition(ZOI) diameter were measured in mm and compared with the standards Gentamicin and Fluconazole for the respective plates[18].

### Minimum Inhibitory Concentration (MIC)

Fungal extract of *A. purpureonitens* was tested for MIC against all microbes used for agar well diffusion[7]. MHB was used as the growth medium for this test. The test microorganisms were cultured in MHB, maintaining the growth at 0.5 MacFarland standard. The analysis was carried out in a 96-well plate, with every test executed in triplicate. 100 microliters of broth was introduced to each well, succeeded by 100 µL of fungal extract at a concentration of 1000µg/mL in the uppermost row of the plate. Pipetting was done 5-6 times to ensure thorough mixing. Serial dilution was then performed for the dilutions 1000 µg/mL to 7.81 µg/mL, with 100 µL transferred for each dilution. Test microorganism suspensions were diluted 20 times using the broth. Subsequently, 10 µL of this bacterial suspension was pipetted into each well, ensuring proper mixing. The plate was sealed and incubated overnight at 37°C. Then, 20 µL of 0.125% TTC (2,3,5-triphenyltetrazolium chloride) dissolved in sterile distilled water was added to each well and incubated for another 2-4 hours.Wells containing viable bacteria turned red with biofilm formation, while those where bacterial growth was inhibited remained colorless, and there was no biofilm formation. The lowest concentration at which color change is not seen is taken as MIC. Broth with bacteria but without any extract served as the control[19].

### Dual Culture Plate Antagonism Assay

A dual culture assay was performed to assess antagonistic interactions between the endophyte *A. purpureonitens* and fungal plant pathogens *Fusarium oxysporum*(815) and *Pythium spp.*[20]. The pathogens were obtained from Nepal Agricultural Research Council (NARC), Khumaltar, Nepal. One-week-old cultures in PDA were used for the assay. 6 mm diameter mycelium disks of the fungal endophyte and the pathogen were placed opposite to each other, symmetrical distance (1-2 cm) away from the petriplate edge. For the control plates, only the phytopathogens were placed at the same position on the respective plates. The PDA plates were incubated at 27±1°C for up to 13□days. Three replicates of each culture combination were carried out.

The growth percentage inhibition of pathogen was calculated as:

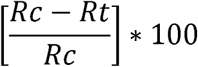

where Rt is the radial growth of the phytopathogen mycelial in the test plates, and Rc is the radial growth of the phytopathogen mycelial in the control plate. The results were displayed as means of the replicates ± standard deviation (SD).

### DPPH(2,2-diphenyl-1-picrylhydrazyl) radical scavenging activity

The DPPH scavenging assay was performed to get the antioxidant capability of the fungal extract[21]. The experiment was conducted over a concentration range of 12.5 to 500 µg/mL, with L-Ascorbic acid in the same concentrations serving as the positive standard[22]. A solution of DPPH was made by dissolving 0.04 mg/mL of DPPH in methanol. 50 µL of the blank(methanol and DPPH) and fungal extract sample of different concentrations was mixed in 96 well plates with 150 µL of the DPPH reagent (1:3 ratio) in each well, respectively. The experiment was carried out in triplicate. The mixtures were then allowed to stay at room temperature in the dark for 30 minutes. After incubation, the absorbance at 517nm was taken in the BioTek Epoch Microplate reader.

The Percentage Inhibition (I%) of L-Ascorbic acid(standard) and the fungal extract, was calculated using the formula:

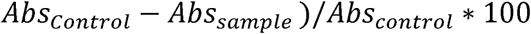

Where Abs_sample_ is the absorbance of the test samples, and Abs_control_ is the absorbance of the control.

A calibration curve was created between the percentage inhibition (I%) and the sample concentrations,then inhibition concentration (IC_50_) was subsequently determined.

### ABTS (2,2**′**-azinobis-(3-ethylbenzthiazolin-6-sulfonic acid)) assay

ABTS assay was carried out following the protocol described by[23, 24] with slight modifications. ABTS radical cations were produced by mixing 2.45 mM of potassium persulfate and 7mM ABTS solution in deionised water. This solution was stored at room temperature in the dark for 12-16 hours. The absorbance of this mixture was taken at 734nm and was set to 0.7 ± 0.02 using a UV-Vis Spectrophotometer, adding methanol for dilution. Dilutions ranging from 12.5 to 500µg/ml were made for both the fungal extract as well as L-Ascorbic acid solution, which was taken as the standard. The sample and ABTS mixture were added in a 1:3 ratio in a 96-well plate, and each sample was triplicated. After a 45-minute incubation in the dark, the absorbances were measured using a BioTek Epoch Microplate reader at 734nm. The absorbance was then plotted as a standard graph. The scavenging activity percentage (%) of the standard as well as the fungal extracts were calculated using the given formula:

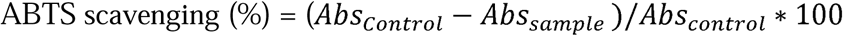

Where Abs_sample_ is the absorbance of the test samples and Abs_control_ is the absorbance of the Control.

The IC□□ value (50% inhibition of ABTS radicals) was determined from the calibration curves generated using different concentrations of the test samples.

### Total Phenolic Content(TFC)

Folin-Ciocalteu reagent Colourimetric Assay was conducted using 96-well plate to find out the TPCof the *A. pupureonitens* extract[25]. Gallic acid was utilized as a standard to measure TPC in the fungal extract. Gallic acid solutions of concentrations from 160, 80, 40, 20, 10, 5 µg/mL were prepared to obtain a calibration curve. A 1 mg/mL fungal crude extract solution was prepared in methanol. 20 µL of standard solution or sample was added to each well, 100 µL Folin-Ciocalteu reagent (prepared in a 1:10 ratio with distilled water) was added to the wells and mixed. The mixture was kept for 10 minutes in the dark then 80 μL sodium

carbonate(Na□CO□) solution (75 g/L) was added, and it was kept in the dark for 1 hour. The experiment was conducted in triplicate using a 96-well plate. After the completion of incubation, the absorbance was taken on BioTek Epoch Microplate reader at wavelength of 750 nm, with methanol as blank.

A standard curve for gallic acid was created, and the concentration of phenolic compounds in the fungal extract was determined using the given formula:

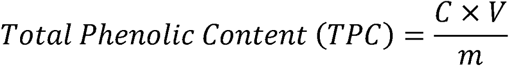

Where “C” is the concentration of the extract (mg/ml) while “V”is the volume of the fungal extract used in mL, and “m” is the weight of the dry extract per ml in g.

The results were reported as micrograms of gallic acid equivalent per each milligram of the fungal extract.

### Total Flavonoid Content(TFC)

The TFC of the *A. purpureonitens* was obtained by the aluminum chloride colorimetric assay with sodium acetate, based on the method described by (Sari et al., 2023) with slight modifications. Quercetin was employed as the reference standard for the preparation of the calibration curve. Standard quercetin solutions were prepared at concentrations of 5µg/mL to 160 µg/mL. For the assay, a stock solution of the fungal extract was made at a concentration of 1 mg/mL. 100 µL of the extract solution was transferred into a 2 mL microcentrifuge tube and mixed thoroughly with 400 µL of methanol. Subsequently, 100 µL of 10% aluminum chloride (AlCl□) solution was added, followed by the adding of 1 M sodium acetate(100µL). The reaction mixture was then incubated in the dark for 45 min for color development. Then, reaction mixture(200 µL) was dispensed into each well of a 96-well microplate, and the absorbance was recorded at 510nm by a BioTek Epoch microplate reader. Measurements were taken in triplicate. The same procedure was repeated for quercetin standard, and a calibration curve was made to quantify the flavonoid content. The total flavonoid content of the sample was presented as milligrams(mg) of quercetin equivalents (QE) per gram of dry extract according to the following equation

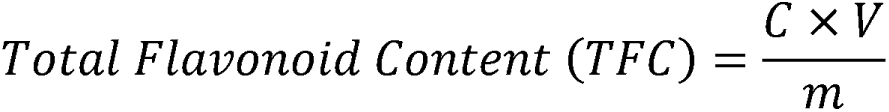

where *C* represents the concentration obtained from the quercetin calibration curve, *V* denotes the volume of the extract used in the assay, and *m* refers to the mass of the dried fungal extract.

### Anticancer Activity

Anticancer activity of the fungal extract was determined applying MTT (3-(4,5-dimethylthiazol-2-yl)-2,5-diphenyltetrazolium bromide) colorimetric assay using human cancer cell lines in 96-well plates following protocols by [27] with slight modifications.. Fungal extract cytotoxicity was tested on the Human cervical adenocarcinoma(HeLa) cell line (acquired from PGIMER, India) and MCF-7 breast cancer cell line (acquired from Shikhar Biotech Pvt. Ltd.,Nepal).

The experiment was performed following the protocol of. The cell lines were culutred in Dulbecco’s Minimum Essential Medium (DMEM) (supplemented with 10% FBS, 200□mM L-glutamine, 7.5□% sodium bicarbonate,100 mM sodium pyruvate1% and Penicillin-Streptomycin). The cells were incubated at 37°C in 5% CO□ conditions until 80% confluency was achieved, after which they were separated using 0.5% trypsin-EDTA. After 80% confluency, the cells were trypsinised, suspended in fresh media and counted using a hemocytometer. The cells were seeded in 96-well plates at 3.5□×□10^3^ cells/well and 4□×□10^3^ cells/well for HeLa and MCF-7 cells, respectively. The seeded cells were incubated for 16 hours in a 5% CO incubator at 37°C and 95% RH.

After 16 hours, as the cells had attached to the plates, the existing media from the 96-well plate was removed, and they were treated with different final concentrations (15-240 µg/mL) of the fungal extract prepared using fresh media. All the treatments were done in triplicate, and 0.1% DMSO was taken as the negative control. The plate was further incubated for 72 hours in the CO□ incubator. Then, 10 µL of 5 mg/ml MTT dye solution (HiMedia, India) in 1X PBS (Thermo Fisher Scientific Inc., USA) was introduced to the wells. After a 3-hour incubation, MTT was reduced to purple water-insoluble formazan crystals, indicating the viable cells. The media was discarded from the wells and the formazan product was dissolved by adding 150 µL DMSO. The microplate reader (MULTISKAN GO Thermo Scientific) was operated at 570 nm wavelength for absorbance reading.

The cytotoxicity results were taken as IC_50_ (50% growth inhibition concentration for the cell lines.

The viability of cell was calculated by:

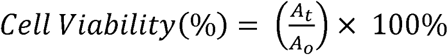

Where ‘A_o_’ is cell line absorbance without fungal extract addition, and ‘A_t_’ is the absorbance with fungal extract addition.

### Liquid Chromatography (LC-MS/MS)

A high-resolution LC–MS/MS evaluation was conducted utilizing quadrupole time-of-flight (QTOF) mass spectrometer (Model G6550A, Agilent Technologies, USA) fitted with a Dual AJSa source of electrospray ionization (ESI) functioning in negative and positive ion modes examined by Sophisticated Analytical Instrumentation Facility (SAIF), Mumbai. The system was governed byAgilent MassHunter Workstation Software was used, and data processing was carried out with Agilent MassHunter.Qualitative Study B.06. Chromatographic separation (Wubshet et al., 2013) was performed on a Hypersil GOLD C18 column (100 × 2.1 mm, 3 µm particle size) kept at 40 °C. The mobile phase comprised solvent A (0.1% formic acid in Milli-Q water) and solvent B (acetonitrile). The gradient program was set as: 0–1 min (5% B), 1–25 min (5–100% B), 25–30 min (100% B), 30–31 min (100–5% B), and 31–35 min (5% B). The volume of injection was 5 µL, and the flow rate was 0.3 mL min ¹. The mass spectrometer was operated under the subsequent optimized settings: capillary voltage (VCap) 3.5 kV, skimmer 65 V, fragmentor voltage 175 V, nozzle voltage 1.0 kV, and octopole RF peak 750 V. The nebulizer pressure was set at 35 psig and temperature of a drying gas was 250 °C (13 L min□¹) with a sheath gas temperature of 300 °C (11 L min□¹). Data were collected in AutoMS² mode over the m/z range 120–1200 with a scanning speed of 1 spectrum per second[28].

### GNPS-Based Spectral Annotation, MASST Searches, and Reverse Metabolomics

The raw LC–MS/MS data were converted to the mzML format and centroided using MSConvert prior to downstream analysis. The processed mzML files were uploaded to the Global Natural Products Social Molecular Networking (GNPS) platform for molecular networking and spectral annotation [29]. In parallel, the centroided mzML files were analyzed using SIRIUS for molecular formula determination and metabolite annotation/structure elucidation based on MS/MS fragmentation patterns and database searching [30]. SIRIUS integrates molecular formula identification with structure annotation tools, including CSI, enabling complementary annotation of LC–MS/MS features [30].

Molecular networking was performed using the GNPS Feature-Based Molecular Networking workflow (Task ID: b92743f54828431bbba68dce36a18014). The precursor ion mass tolerance was set to 0.02 Da, while the fragment ion mass tolerance was set to 0.05 Da. Molecular network edges were retained only when the cosine similarity score exceeded 0.70 with a minimum of six matched fragment ions. For metabolite annotation, all MS/MS spectra were searched against the complete set of GNPS spectral libraries, including experimental reference libraries, propagated libraries, and libraries containing synthetic compounds and conjugated metabolites. Spectral matches were accepted only when they satisfied a cosine similarity score of ≥0.70 with at least six matched fragment ions. All putative annotations were subsequently subjected to manual validation by comparing the experimental and reference MS/MS spectra using mirror plots to confirm fragmentation pattern agreement and improve annotation confidence. To investigate the distribution of the annotated metabolites across publicly available mass spectrometry datasets, the validated spectra were queried using both MicrobeMASST [31] and PlantMASST[32] within the GNPS ecosystem. In addition, reverse metabolomics [33] searches were performed to identify public datasets containing matching MS/MS spectra for the annotated metabolites. Dataset metadata associated with the spectral matches were examined to determine their biological origin where available. Because a substantial number of public datasets lack comprehensive metadata, many spectral matches could not be assigned to a specific organism, sample type, or geographic source. Nevertheless, all matching datasets identified through the reverse metabolomics workflow were retained and reported in the Supplementary Information.

## Results

### Isolation and molecular identification of *A. purpureonitens* from *Taxus wallichiana*

We characterized endophytic fungi associated with *Taxus wallichiana* using an integrated workflow encompassing isolation, molecular identification, metabolite profiling and functional screening (Fig.1). Endophytes were recovered from surface-sterilized leaves purified by subculturing, and taxonomically assigned by PCR amplification and sequencing of the ITS region followed by sequence-based identification. Using this pipeline, isolated T2B was first examined by colony morphology and microscopy (Fig. 1b) and subsequently identified by ITS sequencing as *A. purpureonitens*. The ITS sequence of T2B matched *A. purpureonitens* with 100% query coverage and 100% identity and has been deposited in GenBank under accession PV648432. Phylogenetic placement (Fig. 1c) was performed on Phylogeny.fr v2.0 [34].

**Figure 1.**
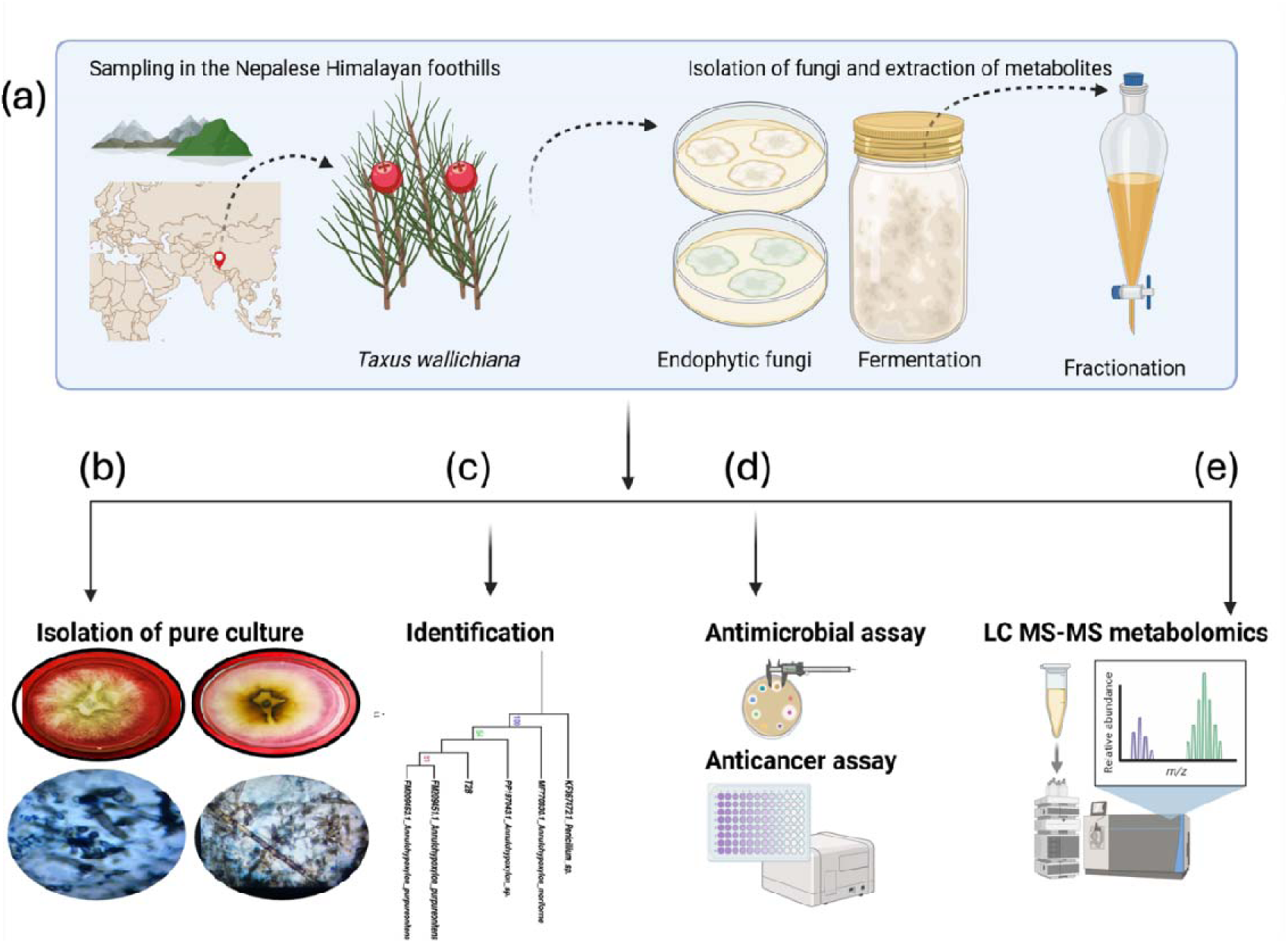
Graphical representation of study design, identification, bioactivity screening and metabolomics of the endophytic fungus. (a) Overview of the workflow for isolating, identifying and characterizing endophytic fungi from *Taxus wallichiana*. Healthy tissues were surface-sterilized and plated on Modified Melin-Norkrans medium (MMN)to isolate endophytes. Pure cultures were obtained by subculturing and identified by genomic DNA extraction, PCR amplification and sequencing of the ITS region, followed by sequence-based taxonomic assignment. Extracts were prepared for bioactivity screening and LC–MS/MS-based metabolite profiling. **(b)** Colony morphology (obverse and reverse) and microscopic features of *Annulohypoxylon purpureonitens* isolate T2B grown on Potato Dextrose Agar (PDA). Colonies were circular with dense cottony–velvety mycelium; the obverse was whitish to pale cream centrally with a subtle star-like pattern and a pale greenish-white to light olive periphery. The reverse showed central yellow–brown pigmentation without pigment diffusion into the agar. Microscopy revealed septate hyphae, predominantly hyaline to light brown, with variable thickness and occasional pigmented segments. **(c)** ITS-based phylogenetic tree showing the placement of isolate T2B among related taxa retrieved from GenBank (BLASTn). The tree was generated using Phylogeny.fr (v2.0) with *Penicillium sp*. as the outgroup; bootstrap values are based on 1,000 replicates. **(d)** Bioactivity screening of the *A. purpureonitens* extracellular extract, including antibacterial activity against human pathogens, antifungal activity against phytopathogens (dual-culture antagonism) and cytotoxicity against cancer cell lines. **(e)** LC–MS/MS-based metabolomics workflow used to annotate and contextualize secondary metabolites detected in the extract, including data acquisition in positive and negative ionization modes, computational annotation (SIRIUS) and GNPS-based molecular networking with MicrobeMASST and PlantMASST to support metabolite prioritization.). This figure (section a,d,e) is created in BioRender [35].

### Bioactivity screening of *Annulohypoxylon purpureonitens* extract Antibacterial activity against human pathogens

The antimicrobial activity of the crude extract of *A. purpureonitens* was evaluated against four bacterial pathogens and one yeast strain using agar well diffusion and minimum inhibitory concentration (MIC) assays. All experiments were performed in triplicate, and the results are expressed as mean ± standard deviation (Table 2). The crude extract exhibited antimicrobial activity against all tested microorganisms, although the degree of susceptibility varied among the strains (Fig. 2a). Among the bacterial strains, *S. aureus* exhibited the largest inhibition zone (15.67 ± 0.58 mm), followed by *E. faecalis* (13.33 ± 1.53 mm), *K. pneumoniae* (10.67 ± 0.58 mm), and *E. coli* (9.33 ± 0.58 mm). The extract also inhibited the growth of *C. albicans*, producing a zone of inhibition of 9.33 ± 0.58 mm. The reference antibiotic gentamicin (10 µg) produced substantially larger inhibition zones against all bacterial strains, ranging from 19.67 ± 1.53 to 32.33 ± 0.58 mm, whereas fluconazole (24 µg) produced a zone of inhibition of 20.67 ± 1.53 mm against C. albicans. The lowest MIC value (500 µg/mL) was recorded for the Gram-positive bacteria *E. faecalis* (ATCC 19433) and S*. aureus* (ATCC 12600), indicating greater susceptibility to the fungal extract. In contrast, the Gram-negative bacteria *K. pneumoniae* (ATCC 13883) and *E. coli* (ATCC 19433), as well as the yeast *C. albicans* (ATCC 10231), exhibited higher MIC values of 1000 µg/mL. Although the activity of the *A. purpureonitens* extract was lower than that of the standard antimicrobial agents, its consistent inhibitory effects highlight its potential as a source of bioactive antimicrobial compounds

**Fig. 2.**
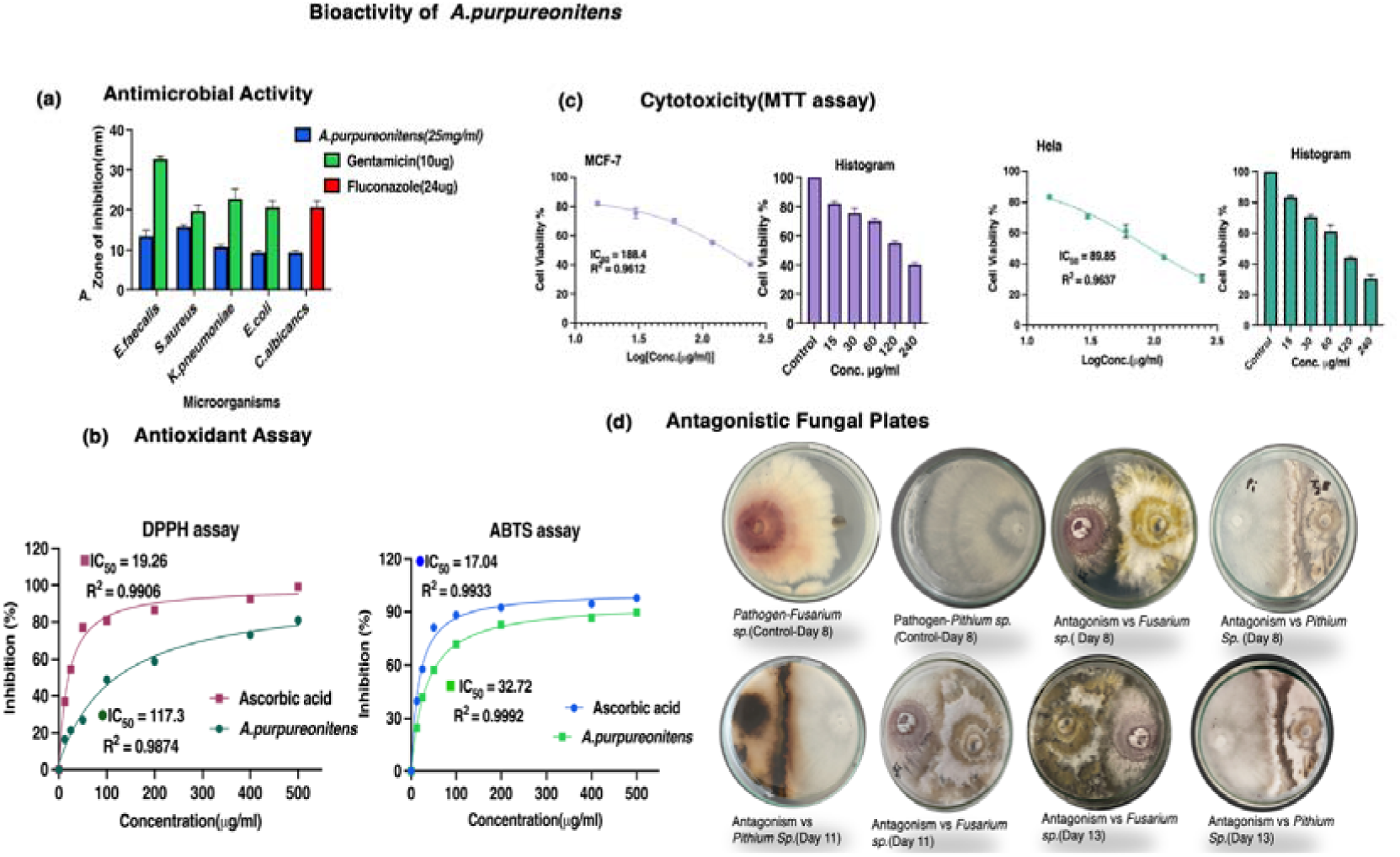
Bioactivity profile of *A. purpureonitens* extracellular extract: **(a)** Antimicrobial activity represented by zones of inhibition (mm) against target human pathogens across treatment concentrations compared to control. **(b)**Antioxidant activity evaluated via DPPH and ABTS free-radical scavenging assay with standard ascorbic acid. **(c)**In vitro cytotoxic activity illustrates cell inhibition percentages across cell lines (HeLa and MCF-7) with corresponding IC50 values (μg/mL). Data are expressed as mean ± SEM(n=3) **(d)** Representative dual-culture antagonistic assays against fungal phytopathogens (*Fusarium sp.*and *Pithium sp.*) compared to pathogen-only controls. Data are expressed as mean ± SD (n=3).

**Figure 3.**
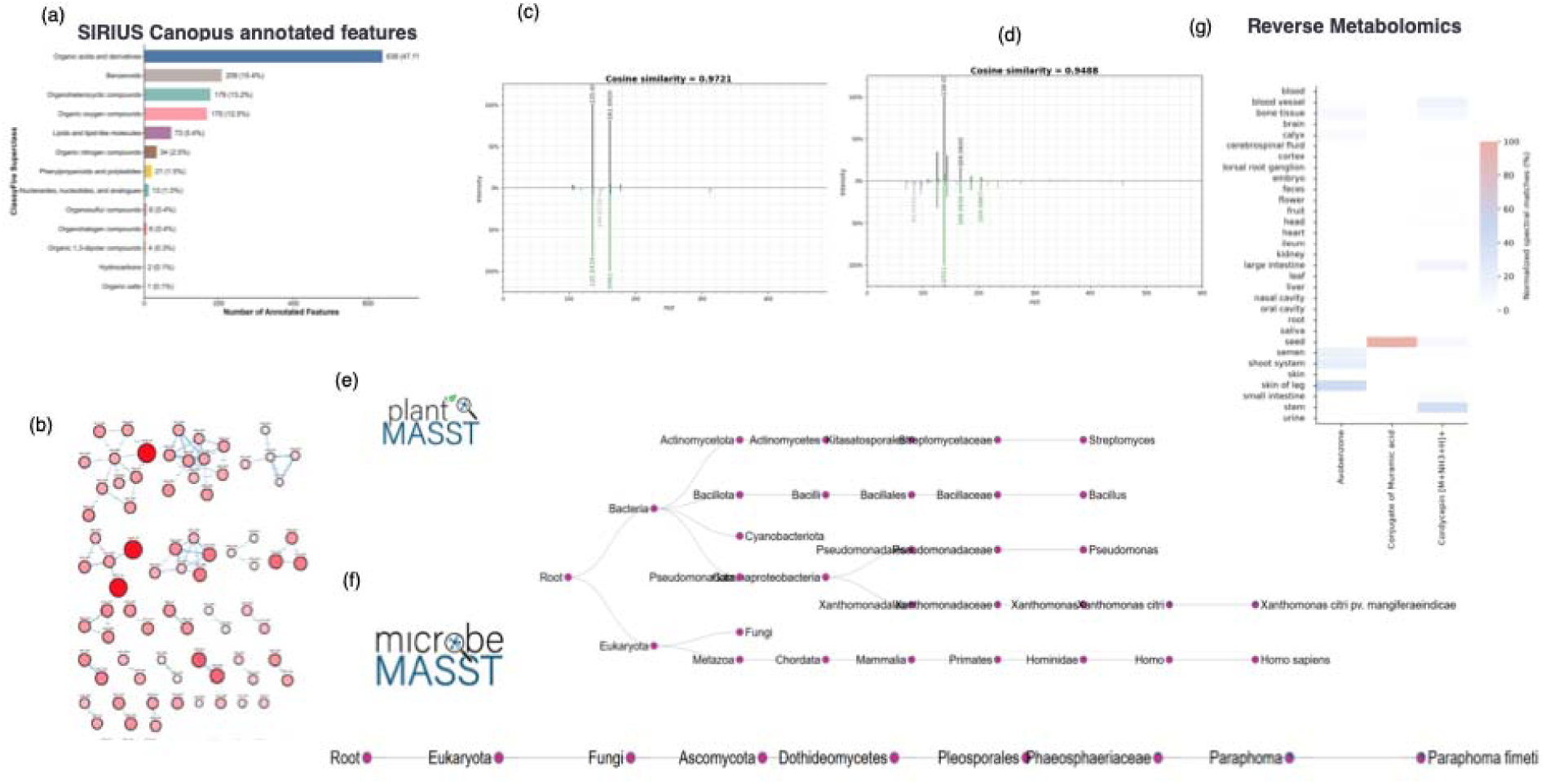
Reverse metabolomics and MASST analysis reveal the widespread occurrence of annotated metabolites across public mass spectrometry datasets. *A. purpureonitens* extract. **(a)** Distribution of annotated molecular features identified using the SIRIUS/Canopus workflow according to predicted chemical classes of positive ionization mode and negative ionization mode (Fig.S1)**. (b)** GNPS molecular networking was performed using all available experimental and propagated spectral libraries, including synthetic and conjugated compound libraries**. (c–d)** Candidate metabolite annotations were manually validated by comparison of experimental and reference MS/MS spectra using mirror plots. **(e)** PlantMASST searches were performed to investigate their occurrence in plant-associated datasets **(f))** MicrobeMASST searches identified matching spectra for the annotated metabolites across diverse microbial datasets.. **(g)** Reverse metabolomics analysis further demonstrated that the muramic acid conjugate and cordycepin conjugate are present in thousands of publicly available mass spectrometry datasets, although many lack sufficient metadata for taxonomic or environmental assignment. Representative microbial taxa associated with matching datasets include Actinomycetes, *Bacillus spp., Pseudomonas spp., Streptomyces sp*., *Salinispora, Nocardia, Alternaria, Mortierella alpina,* and *Paraphoma fimeti*

**Table 2.**
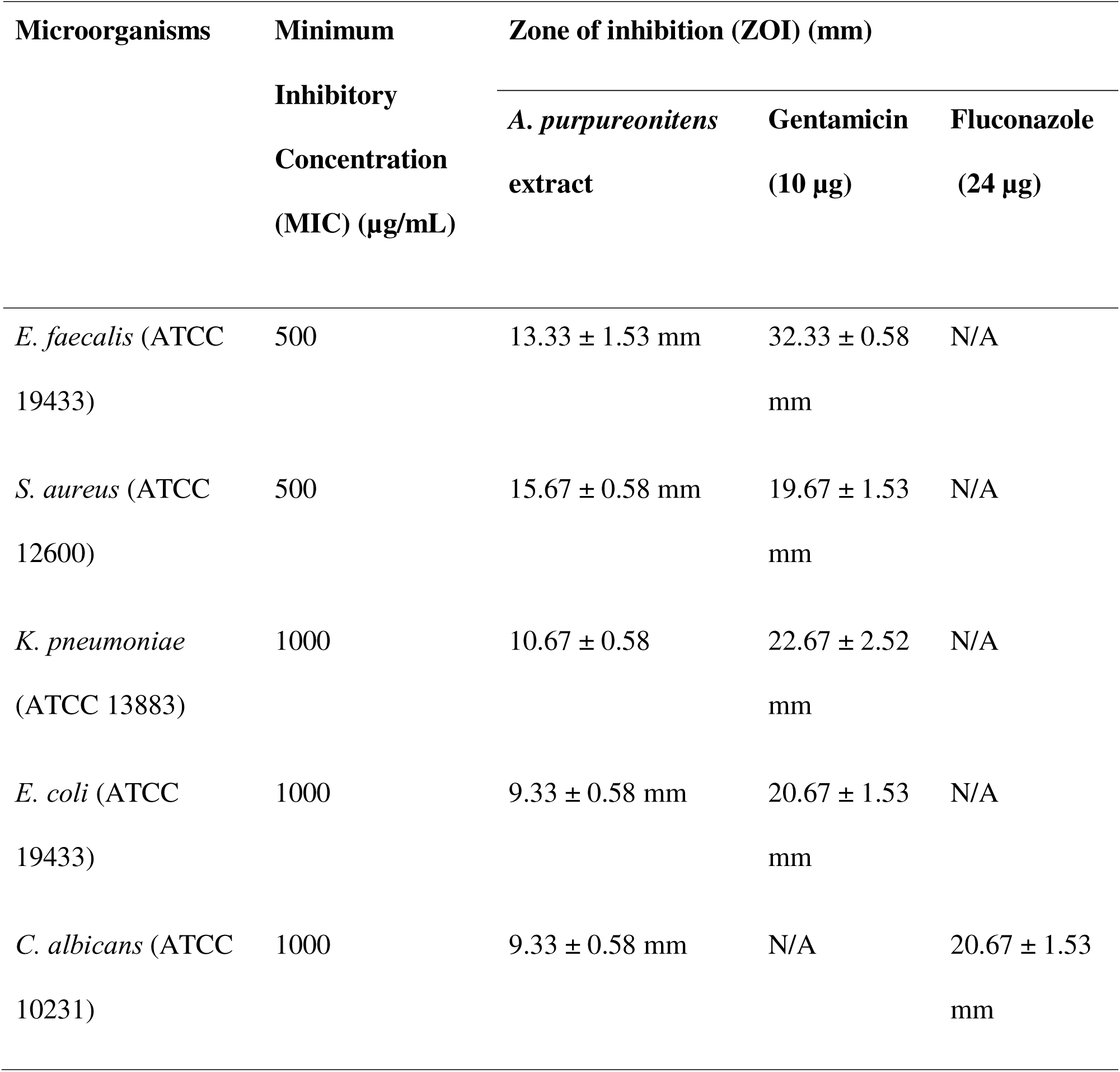
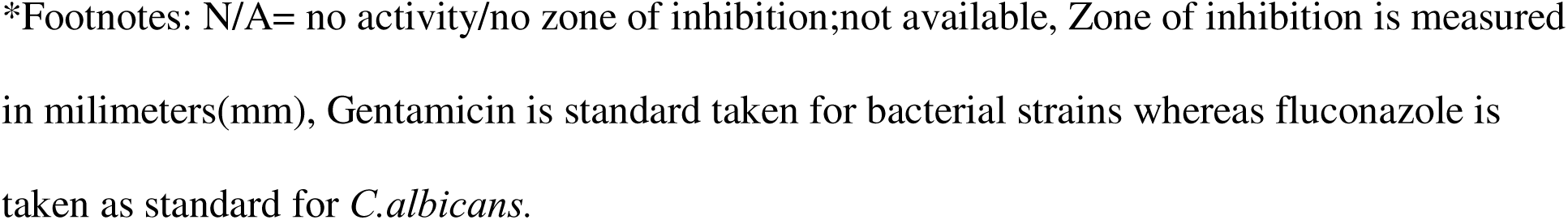
Antimicrobial activity of the *A.purpureonitens* extract against selected bacterial and fungal pathogens, expressed as zone of inhibition (mean ± SD), compared with gentamicin and fluconazole, together with the minimum inhibitory concentration (MIC) of the extract.

| Microorganisms | Minimum Inhibitory Concentration (MIC) ($\mu\text{g/mL}$) | Zone of inhibition (ZOI) (mm) | | |
| --- | --- | --- | --- | --- |
| | | *A. purpureonitens* extract | Gentamicin (10 $\mu\text{g}$) | Fluconazole (24 $\mu\text{g}$) |
| *E. faecalis* (ATCC 19433) | 500 | 13.33 ± 1.53 mm | 32.33 ± 0.58 mm | N/A |
| *S. aureus* (ATCC 12600) | 500 | 15.67 ± 0.58 mm | 19.67 ± 1.53 mm | N/A |
| *K. pneumoniae* (ATCC 13883) | 1000 | 10.67 ± 0.58 | 22.67 ± 2.52 mm | N/A |
| *E. coli* (ATCC 19433) | 1000 | 9.33 ± 0.58 mm | 20.67 ± 1.53 mm | N/A |
| *C. albicans* (ATCC 10231) | 1000 | 9.33 ± 0.58 mm | N/A | 20.67 ± 1.53 mm |

\*Footnotes: N/A= no activity/no zone of inhibition;not available, Zone of inhibition is measured in millimeters(mm), Gentamicin is standard taken for bacterial strains whereas fluconazole is taken as standard for *C.albicans*.

### Dual culture antagonism assay

The dual culture antagonism assay displays the inhibitory effect of the endophytic fungi on target plant pathogens in the dual culture plate, showing matrix competition type effect as described by [36]. Specifically, the growth resembles ‘Mutual slight inhibition’ type growth in the *A. purpureonitens* vs *Pythium* plate, and ‘Growth around’ type growth in the *A. purpureonitens* vs *Fusarium oxysporum* plate [37] The endophyte showed similar inhibition rates when confronted with both fungal pathogens. The test plates (Fig.2d) evaluated on the eighth day of culture showed that the endophyte exhibited similar inhibition rates (Mean ± S.D.) when confronted with both fungal pathogens. For *Fusarium oxysporum,* the growth inhibition rate was found to be 25.35 ± 10.28%. While for *Pythium spp.,* the growth inhibition rate was slightly higher at 29.17 ± 8.78%. Thus, the endophyte demonstrated significant antagonistic influence towards the test pathogens.

### Antioxidant capacity and radical-scavenging activity (DPPH and ABTS)

The antioxidant potential of the *A. purpureonitens* crude extract was systematically evaluated using two distinct radical scavenging mechanisms: the DPPH and ABTS assays, with ascorbic acid serving as the positive control. Both assays revealed a distinct, concentration-dependent increase in radical scavenging activity across the tested range of 0 to 500μg/mL (Fig.2b). In the DPPH assay, ascorbic acid exhibited rapid scavenging kinetics with an IC_50_ value of 19.26 μg/mL. The *A. purpureonitens* extract demonstrated a progressive radical-scavenging curve, yielding an IC_50_ value of 117.3 μg/mL and reaching approximately 80.96% inhibition at the highest tested concentration.

Interestingly, the extract showed significantly higher efficacy in the ABTS assay, closely approaching the potency of the standard reference. The reference standard cleared the ABTS^+^ radical with an IC_50_ of 17.04 μg/mL. The *A. purpureonitens* extract followed a highly competitive curve, yielding a remarkably low IC_50_ value of 32.72 μg/mL and stabilizing at over 89.61% inhibition at higher concentrations.

### Total phenolic and flavonoid contents

The quantitative estimation of secondary metabolites revealed notable concentrations of phenolic and flavonoid compounds in the fungal extract. The concentration of TPC in fungal extract was found to be 10.43 ± 1.37 w/w% gallic acid equivalents (GAE) whereas the concentration of TFC in the sample was found to be 2.29 ± 0.02 quercetin equivalent (QE) per gram. The corresponding calibration curve of gallic acid and quercetin are presented in Fig.S3. These concentrations suggest that phenolic compounds constitute a major fraction of the extract’s bioactive secondary metabolites, which likely contribute to the observed antioxidant and antimicrobial activities.

### Cytotoxicity activity against Cancer cell lines

Cytotoxic potential of *A.purpureonitens* extract was analyzed against two human cancer cell lines, MCF-7 and HeLa. The extract demonstrated dose-dependent toxicity in both cell lines across the examined concentration range of 15–240 µg/mL (Fig.2c). A gradual decrease in cell viability was noticed with higher extract concentration, suggesting a relationship dependent on concentration suppressive influence. The IC□□ values were found to be 188.4 µg/mL for the MCF-7 cell line and 89.85µg/mL for the HeLa cell line which suggests slightly higher sensitivity of HeLa cells to the extract compared to MCF-7. Cell viability was reduced to 40.3% in MCF-7 cells and 30.6% in HeLa cells at highest concentrations(240µg/ml) Overall, the results demonstrate notable cytotoxic activity of the extract, with differential sensitivity observed between the two cancer cell lines.

### Occurrence of Annotated Metabolites Across Public Mass Spectrometry Datasets Revealed by MASST and Reverse Metabolomics

Following GNPS-based molecular networking, the MS/MS spectra obtained from the *A. purpureonitens* extract were systematically annotated using multiple complementary approaches. First, molecular networking was performed in GNPS against all available spectral libraries, including both experimental and propagated libraries containing synthetic and conjugated compounds. Candidate annotations were subsequently validated by manual inspection of MS/MS mirror plots to confirm spectral similarity and fragment ion agreement. To investigate the broader occurrence of the annotated metabolites, MicrobeMASST and PlantMASST searches were performed, followed by reverse metabolomics analysis across publicly available mass spectrometry datasets. This analysis revealed that several annotated metabolites, particularly the muramic acid conjugate and the cordycepin conjugate, were repeatedly detected in numerous public datasets, indicating that these metabolites are widely distributed across diverse biological samples. MicrobeMASST analysis showed that spectra corresponding to the cordycepin conjugate were detected in datasets associated with a wide range of microorganisms, including *Actinomycetes, Bacillus amyloliquefaciens, Bacillus subtilis, Bacillus spp., Pseudomonas fluorescens, Pseudomonas donghuensis, Pseudomonas putida, Pseudomonas chlororaphis, Pseudomonas vranovensis, Erwinia endophytica, Nocardia spp., Streptomyces spp., Streptomyces nojiriensis, Salinispora, Mycolicibacterium smegmatis* MC2 155*, Mycobacterium palustre, Bacteroides fragilis, Xanthomonas citri pv. mangiferaeindicae, Polaribacter porphyrae*, members of the *Cyanobacteriota, Mortierella alpina*, *Alternaria*, and other fungal taxa. Similarly, the avobenzone conjugate was also detected across multiple microbial datasets, while the muramic acid conjugate was identified in datasets associated with the fungus *Paraphoma fimeti*. Notably, reverse metabolomics searches identified matches for the muramic acid and cordycepin conjugates (Fig.S2) in thousands of publicly available datasets. However, a substantial proportion of these datasets lack comprehensive metadata, preventing reliable determination of their biological origin, sample type, or geographic source. Nevertheless, the corresponding dataset identifiers and spectral matches are provided in Supplementary Table 1, enabling future exploration as additional metadata become available. The complete list of matching datasets and associated metadata, where available, is provided in Supplementary Figure 1.

## Discussion

This study provides an integrated assessment of the biological activity and chemical space of *A. purpureonitens*, an endophytic fungus isolated from *Taxus wallichiana*. Endophytic fungi are increasingly recognized as reservoirs of structurally diverse secondary metabolites with antimicrobial, antioxidant and cytotoxic activities, partly reflecting their ecological association with plant hosts [38].In this context, the combination of phenotypic screening with LC–MS/MS molecular networking, spectral validation and public-dataset searches provides a framework for connecting biological activity with the chemical space of an underexplored endophyte. Our findings indicate that *A. purpureonitens* produces an extracellular metabolome associated with measurable antibacterial, antifungal, antioxidant and cytotoxic phenotypes.

The antibacterial activity of the crude extract was selective rather than uniformly broad-spectrum. The stronger activity observed against Gram-positive organisms is compatible with differences in cell-envelope architecture between Gram-positive and Gram-negative bacteria, although such differences alone cannot establish a mechanism of action[39]. Importantly, the relatively high MIC values indicate that the activity should be interpreted as a property of a complex crude extract rather than evidence for a highly potent single antimicrobial compound. The observed activity is therefore better viewed as a phenotype supporting further fractionation and chemical characterization. Similarly, the fungal confrontation assay demonstrated antagonistic activity against the tested phytopathogens, but this type of assay reflects the net outcome of a living fungal interaction and cannot distinguish chemical inhibition from nutrient competition, differential growth rates or other ecological interactions. Thus, the observed inhibition provides evidence for antagonistic potential but does not establish the activity of a specific secondary metabolite.

The antioxidant and cytotoxicity assays provide complementary evidence of biological activity. The different responses observed in the DPPH and ABTS assays are consistent with differences in their reaction chemistry and demonstrate why antioxidant capacity should not be inferred from a single assay[40]. The relatively high total phenolic content further supports a possible contribution of reducing or phenolic constituents, although colorimetric TPC and TFC measurements provide class-level estimates and cannot establish the identity or abundance of individual metabolites. The cytotoxicity observed against HeLa and MCF-7 cells should likewise be regarded as preliminary activity of the crude extract. Differences in sensitivity between cell lines are common for fungal extracts and their metabolites, but the concentrations required for activity and the complex composition of the extract preclude assigning the phenotype to individual compounds. Isolation, fractionation and selectivity testing against non-transformed cells will therefore be required before therapeutic relevance can be assessed.

The LC–MS/MS analysis substantially expanded the chemical context of these phenotypes. GNPS molecular networking and spectral-library searching enabled structurally related MS/MS features to be organized and compared with reference spectra, providing an efficient dereplication strategy for complex natural-product extracts [29, 41]. Importantly, the compound assignments in this study should be considered putative Level 2/probable annotations, rather than confirmed metabolite identifications. The annotations were supported by precursor mass, spectral-library similarity and manual MS/MS mirror-plot inspection, but authentic-standard comparison and retention-time matching were not performed. Under the confidence framework proposed by Schymanski et al., such spectral evidence corresponds to probable structures rather than Level 1 confirmed identifications [42]. This distinction is particularly important for structural isomers and conjugated metabolites, which may generate similar fragmentation patterns. Accordingly, the annotated compounds should be regarded as hypotheses for targeted validation rather than definitive chemical identifications.

The combination of molecular networking with MASST and reverse metabolomics provided an additional layer of information beyond conventional library annotation. MASST enables individual MS/MS spectra to be searched across publicly available datasets, thereby revealing where similar spectra have previously been observed [43]. In the present study, repeated spectral matches were observed for several annotated metabolites, particularly the muramic-acid-and cordycepin-related conjugates, across diverse microbial datasets.

The putative annotation of hydroquinidine-, chlorogenic acid, muramic-acid-and cordycepin-related conjugates is noteworthy because their corresponding parent structures have established biological relevance. These metabolites or structurally related compounds have been associated with diverse biological activities, including antioxidant and antimicrobial effects, and may therefore contribute to the observed bioactivity of the extract. However, these annotations should be considered tentative until confirmed using authentic standards and/or complementary spectroscopic analyses. [44–47].

These observations are biologically interesting and could be responsible compound for biol but should not be interpreted as evidence that the corresponding organisms necessarily biosynthesize the exact compounds assigned by spectral annotation. A MASST match establishes similarity between MS/MS spectra, but does not independently establish molecular structure, biosynthetic origin or ecological function. Instead, the widespread occurrence of matching spectra suggests that these chemical features may belong to a broader microbial chemical space and may represent conserved metabolites, conjugates or transformation products. This provides a useful hypothesis for future targeted investigation.

Several limitations should be considered. First, the biological sampling was relatively small, and the observed chemical profile cannot yet be assumed to represent the broader metabolic diversity of *A. purpureonitens* or the endophytic mycobiome associated with *T. wallichiana*. Secondary-metabolite production can vary substantially among fungal strains and culture conditions; therefore, independent isolates and replicated cultures will be required to establish reproducibility. Second, the LC–MS/MS analysis was primarily exploratory and was not designed as a fully quantitative untargeted metabolomics experiment with an extensive pooled-QC framework. Future studies should incorporate biological and technical replicates, extraction blanks, pooled QC samples, internal standards and repeated QC injections to assess feature reproducibility, background contamination and instrumental drift. Pooled QC samples are increasingly recognized as important for evaluating analytical variation and drift in untargeted LC–MS metabolomics [48], while broader QA/QC frameworks emphasize systematic assessment of sample preparation, instrument performance and data reproducibility [49].

Finally, the metabolite annotations require experimental confirmation. Manual mirror plots increase confidence in spectral similarity but cannot establish definitive molecular structures. Priority features should therefore be validated using authentic standards, including retention-time and MS/MS comparison under matched conditions, with NMR or other orthogonal approaches used where structural ambiguity remains. Current metabolomics reporting recommendations emphasize the importance of clearly distinguishing putative annotations from experimentally confirmed metabolites [50].

## Conclusion

Overall, *A. purpureonitens* represents a chemically and biologically interesting, but still underexplored, endophytic fungus associated with *T. wallichiana*. The integration of bioactivity screening, GNPS molecular networking, manual spectral validation and MASST-based reverse metabolomics moves the analysis beyond simple compound-list generation by placing candidate metabolites within a broader microbial chemical context. The next step is targeted isolation and structural validation of the highest-priority features, followed by quantitative and mechanism-oriented studies to determine which metabolites, if any, contribute directly to the observed biological phenotypes.

## Acknowledgements

We would like to acknowledge the Department of Biotechnology, Kathmandu University, Nepal, for providing the research facilities and workspace to conduct this study. We are sincerely thankful to the Swedish University of Agricultural Sciences, Sweden for funding the project.

## Statements and Declarations

### Author contributions

Dhruva Prasad Gauchan: Experiment design, manuscript design and supervision; Rosario Garcia Gill: Research funding, supervision, manuscript editing, Heriberto Vélez: manuscript preparation and supervision; Tara Shrestha: experiment conduction, manuscript writing and data analysis; Santosh Lamichhane: Data analysis and manuscript writing, supervision, editing; Anjela: Manuscript preparation and experiment conduction; Samriddhi Bhochhibhoya: experiment and manuscript writing

### Electronic Supplementary material

Below is the link to electronic supplementary material

### ESM1.pdf

**Supplementary figure 1: Fig. S1** ClassyFire chemical superclass distribution of SIRIUS/Canopus-annotated features from the *A. purpureonitens* extract of negative ionization mode. **Fig. S2** The complete list of matching datasets of conjugate of Muramic acid and Cordycepin. **Fig. S3** Standard curves were generated using simple linear regression. (A) Gallic acid curve for total phenolic content (TPC) and (B) Quercetin for total flavonoid content (TFC). **ESM2.xlxs**

**Supplemtary table 1:** Table S1 and S2 contains spectral matches from GNPS annotion to all availbale datasets in both positive and negative ionization modes.

## Data Availability

The fungus is stored in Department of Biotechnology, Kathmandu University. The mass spectrometry data generated in this study are publicly available through the MassIVE repository under accession number MSV000102749. The DNA sequence generated is submitted to Genebank under the accession number PV648432 and are available at the following url: https://www.ncbi.nlm.nih.gov/nuccore/2978463667

