## Supplementary figure 1 for "Uncovering bioactive metabolites from the *Taxus wallichiana* endophyte *Annulohypoxylon purpureonitens* using reverse metabolomics"

### Bioactivity and reverse metabolomics of an endophytic fungus from Taxus wallichiana

| **Contents** | **Page no.** |
|---|---|
| **Fig. S1** ClassyFire chemical superclass distribution of SIRIUS/Canopus-annotated features from the *A. purpureonitens* extract of negative ionization mode. | 2 |
| **Fig. S2** The complete list of matching datasets of conjugate of Muramic acid and Cordycepin | 3 |
| **Fig. S3** Standard curves were generated using simple linear regression. (A) Gallic acid curve for total phenolic content(TPC) and (B) Quercetin for total flavonoid content(TFC) | 4 |

Corresponding Author's Email:

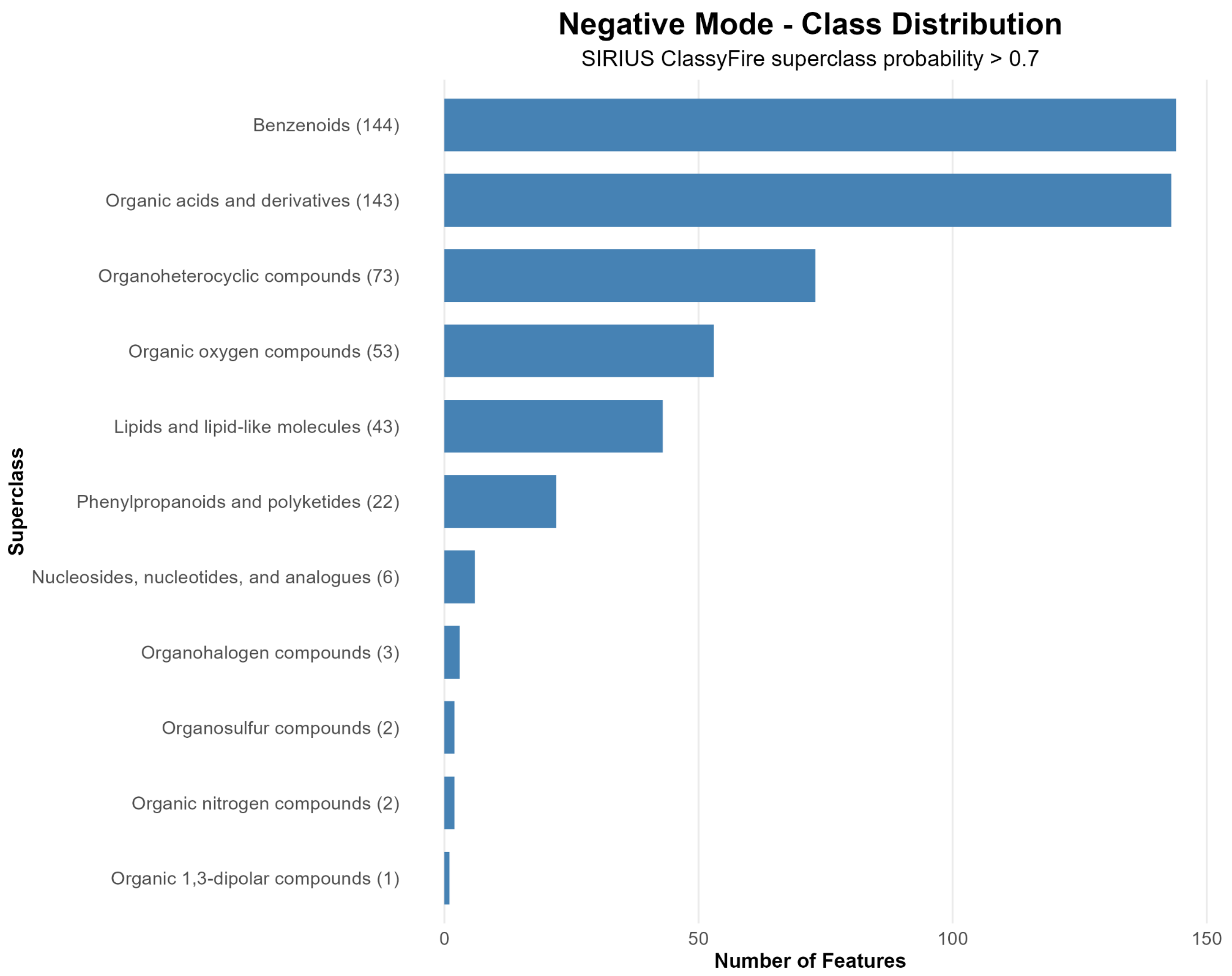


**Fig. S1** ClassyFire chemical superclass distribution of SIRIUS/Canopus-annotated features from the *A. purpureonitens* extract of negative ionization mode.

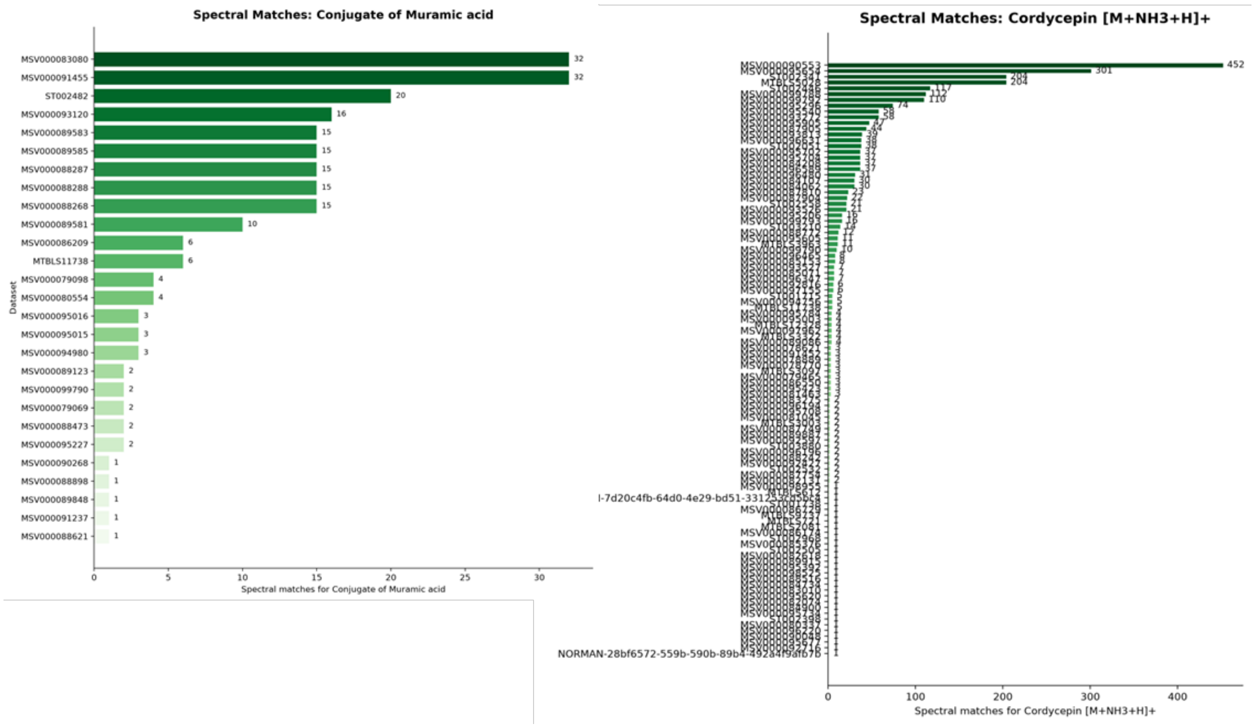


**Fig. S2** The complete list of matching datasets of conjugate of Muramic acid and Cordycepin

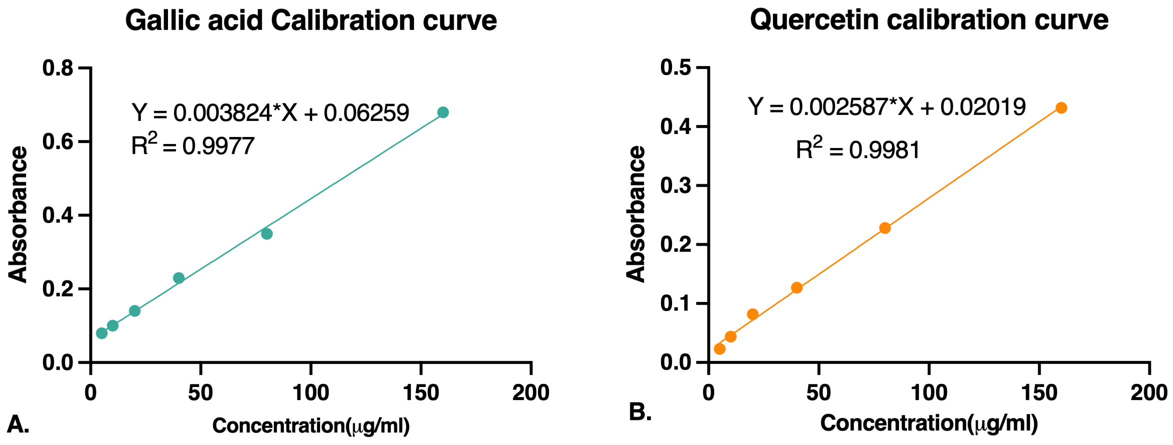


**Fig. S3** Standard curves were generated using simple linear regression. (A) Gallic acid curve for total phenolic content(TPC) and (B) Quercetin for total flavonoid content(TFC)
